# Assessing the variation within the oral microbiome of healthy adults

**DOI:** 10.1101/2020.05.07.083634

**Authors:** Jacob T. Nearing, Vanessa DeClercq, Johan Van Limbergen, Morgan G.I. Langille

## Abstract

Over 1000 different species of microbes have been found to live within the human oral cavity where they play important roles in maintaining both oral and systemic health. Several studies have identified the core members of this microbial community, however, the factors that determine oral microbiome composition are not well understood. In this study we exam the salivary oral microbiome of 1049 Atlantic Canadians using 16S rRNA gene sequencing in order to determine which dietary, lifestyle, and anthropometric features play a role in shaping microbial community composition. Features that were identified as being significantly associated with overall composition were then additionally examined for genera and amplicon sequence variants that were associated with these features. Several associations were replicated in an additional secondary validation dataset. Overall, we found that several anthropometric measurements including waist hip ratio, height, and fat free mass, as well as age and sex, were associated with oral microbiome composition in both our exploratory and validation cohorts. We were unable to validate dietary impacts on the oral microbiome but did find evidence to suggest potential contributions from factors such as the number of vegetable and refined grain servings an individual consumes. Interestingly, each one of these factors on their own were associated with only minor shifts in the oral microbiome suggesting that future biomarker identification for several diseases associated with the oral microbiome may be undertaken without the worry of confounding factors obscuring biological signal.

**Importance:** The human oral cavity is inhabited by a diverse community of microbes known as the human oral microbiome. These microbes play a role in maintaining both oral and systemic health and as such have been proposed to be useful biomarkers of disease. However, to identify these biomarkers, we first need to determine the composition and variation of the healthy oral microbiome. Within this report we investigate the oral microbiome of 1049 healthy individuals to determine which genera and amplicon sequence variants are commonly found between individual oral microbiomes. We then further investigate how lifestyle, anthropometric, and dietary choices impact overall microbiome composition. Interestingly, the results from this investigation showed that while many features were significantly associated with oral microbiome composition no single biological factor explained a variation larger than 2%. These results indicate that future work on biomarker detection may be encourage by the lack of strong confounding factors.

## Introduction

The human oral cavity is colonized by numerous bacteria, fungi, viruses and archaea that make a rich microbial community known as the oral microbiome. This microbial community is one of the most diverse sites of microbial growth within the human body being only secondary to the colon (1). To date over 1000 different bacterial species have been found to colonize the oral cavity (2) on various surfaces including the tongue, teeth, cheek, and gingivae (1). These communities of microbes are responsible for various functions that can both maintain and deplete oral health. For example, the presence of biofilms containing bacterial species such as *Streptococcus mutans* and other aciduric bacteria can damage hard dental surfaces and lead to dental caries (3, 4). Furthermore, the oral microbiome is known to play a role in a myriad of other oral diseases including oral cancer (5), periodontitis (6, 7), and gingivitis (8, 9). In addition to well-established associations between oral and cardiac health (10), recent work has also begun to show that the oral microbiome may play a role in the health of other distal sites within the human body. This includes diseases such as colorectal cancer (11, 12), pancreatic cancer (13), prostate cancer (14), atherosclerosis (15) and inflammatory bowel disease (16).

Due to the associations between these diseases and the oral microbiome, its composition has been proposed as a useful biomarker for human health and disease. With this in mind, various studies have attempted to identify core members of the “healthy” oral microbiome (1, 17–20) to help aid in disease detection. These studies have uncovered that, at the genus level, the oral microbiome remains relatively stable between individuals(1, 20) and across multiple geographic locations(18, 21), but at deeper taxonomic resolutions, it can be variable. This variability has indicated that dietary, anthropometric or sociodemographic factors may play a role in shaping the oral microbiome(17, 19, 22–25). Various studies have focused on individual factors that may cause shifts in the oral microbiome such as ethnicity(1, 25), alcohol consumption(26), smoking(27), obesity(28, 29), and dietary patterns(30). However, to date only a small number of studies have looked at the relative contributions of each of these factors to oral microbiome variability in a single cohort. Takeshita et al., examined the oral microbiome of 2343 adults living in Japan using 16S rRNA gene sequencing and identified that higher abundances of *Prevotella*, and *Veillnella* species were associated with old age, higher body mass index (BMI), and poor overall oral health (19). Another study by Renson et al., in adults living in New York city also found that variation in taxonomic abundances could be linked to marital status, ethnicity, education and age (23). Further, work by Belstrøm et al., examined the oral microbiome of 292 Danish individuals with low levels of dental caries and periodontitis using microarrays and found that while socioeconomic status impacted oral microbiome profiles, diet, BMI, age, and sex had no statistical impact on microbial abundances (22). This study, however, was only able to identify the abundances of taxa that had a corresponding probe which, could explain its disagreement with other work. Overall, these studies have indicated that both biological differences such as sex and BMI as well as lifestyle and sociodemographic differences can impact oral microbiome composition.

While these studies have shed light on the variation of the oral microbiome, it is currently unclear to what extent these factors play a role in shaping the oral microbiome of an individual. Without identifying the effect size of each of these factors relative to one another, it is difficult to identify the correct variables that should be controlled for in case-control studies of the oral microbiome. Furthermore, each of these studies have identified different taxa that are impacted by various factors such as sex, BMI and age. This could be due to many factors, including systemic bias introduced via the use of different protocols or differences in the studied cohorts. Therefore, the identification of microbes that are impacted by factors such as sex, BMI, or diet could help identify potential interactions between the oral microbiome, health, and disease.

Herein, we report the variation within the healthy oral microbiome by examining 741 samples from non-smoking healthy individuals living within the Atlantic Provinces of Canada. We then validated our results on a smaller subset of individuals (n=308) from the same cohort (**Sup Fig 1**). The bacterial oral microbiome composition of these individuals was investigated through 16S rRNA gene sequencing from saliva samples provided by each participant. Compositions were then compared with 41 different variables including anthropometric, dietary and sociodemographic factors. In this investigation, we determined which of these factors play a role in shaping the oral microbiome and to what extent these factors can explain the overall oral microbiome composition.

## Methods

### Study design and population

The current study includes the analysis of saliva samples from the Atlantic Partnership for Tomorrow’s Health (PATH) study. Atlantic PATH is part of the Canadian Partnership for Tomorrow’s Health (CanPath) project, a pan-Canadian prospective cohort study examining the influence of environmental, genetic and lifestyle factors on the development of chronic disease (31). The applicable provincial and regional ethics boards approved the study protocol and all participants provided written informed consent prior to participation. The primary inclusion criteria were that participants were aged 30-74 years at time of recruitment, a resident in one of the Atlantic Canadian provinces (Nova Scotia, New Brunswick, Prince Edward Island, and Newfoundland and Labrador). Recruitment and baseline data for all participating regions was collected between 2000 and 2019. Details on participant recruitment and a descriptive cohort profile have been published elsewhere (31). The questionnaire included sociodemographic information, health information, behaviours, environmental factors, and self-reported anthropometric information. Participants also had anthropometric measures (height, weight, waist and hip circumferences, body composition, blood pressure, grip strength, and resting heart rate) and biological samples (blood, urine, saliva, and toenails) collected. Approximately 9000 participants in the Atlantic PATH cohort provided a saliva sample. Participants were instructed to refrain from eating, smoking, or chew gum for at least 30 minutes prior to oral specimen collection. Oral samples (3 ml) were collected in sterile 50 ml conical tubes after rinsing with water. Samples were stored at 4°C and batch shipped on ice to the central processing facility at the QEII Health Sciences Centre in Halifax, Nova Scotia. Samples were processed within 24 hours of collection, aliquoted into cryovials and stored at −80°C until analysis.

The current analysis includes a total of 1214 saliva samples from healthy Atlantic Canadians living within the provinces of Nova Scotia, New Brunswick, and Prince Edward Island. Based on self-reported data, participants were defined as healthy if they had not been diagnosed with any of the following conditions: hypertension, myocardial infarction, stroke, asthma, chronic obstructive pulmonary disease, major depression, diabetes, inflammatory bowel disease, irritable bowel syndrome, chronic bronchitis, emphysema, liver cirrhosis, chronic hepatitis, dermatologic disease (psoriasis and eczema), multiple sclerosis, arthritis, lupus, osteoporosis, and cancer. A total of 165 of these samples were removed due to insufficient sequencing depth and of the remaining 1049, 308 were removed due to incomplete answering of the 41 variables examined in this study. These 308 samples that were removed were then used in validation analysis (details below) to confirm findings within the larger 741 participant cohort.

### Socio-demographic, lifestyle and anthropometric variables

Questionnaires were used to collect socio-demographic and lifestyle variables. Self-reported variables included age, sex, education level, household income, rural/urban, province, dental visits, sleep patterns, alcohol consumption, smoking status, and dietary variables such as food avoidance, the use of specific types of fat/oil and artificial sweeteners, the frequency of dessert, soda drinks, soy/fish sauce, seasoning with salt seasoning, and fast food, as well as servings of vegetables, fruit, juice, whole grains, refined grains, dairy products, eggs, fish, tofu, beans, and nuts/seeds. Anthropometric measures were collected by trained personnel in assessment centres. Waist and hip circumferences were measured using Lufin steel tape. Height was measured by a Seca stadiometer. Height and weight measures were used to calculate body mass index (BMI; weight in kilograms divided by height in meters squared; kg/m^2^). Body weight, fat mass, and fat-free mass were measured using the Tanita bioelectrical impedance device (Tanita BC-418, Tanita Corporation of America Inc., Arlington Heights, Illinois). Table 1 lists all variables that were used for analysis.

**Table 1:**
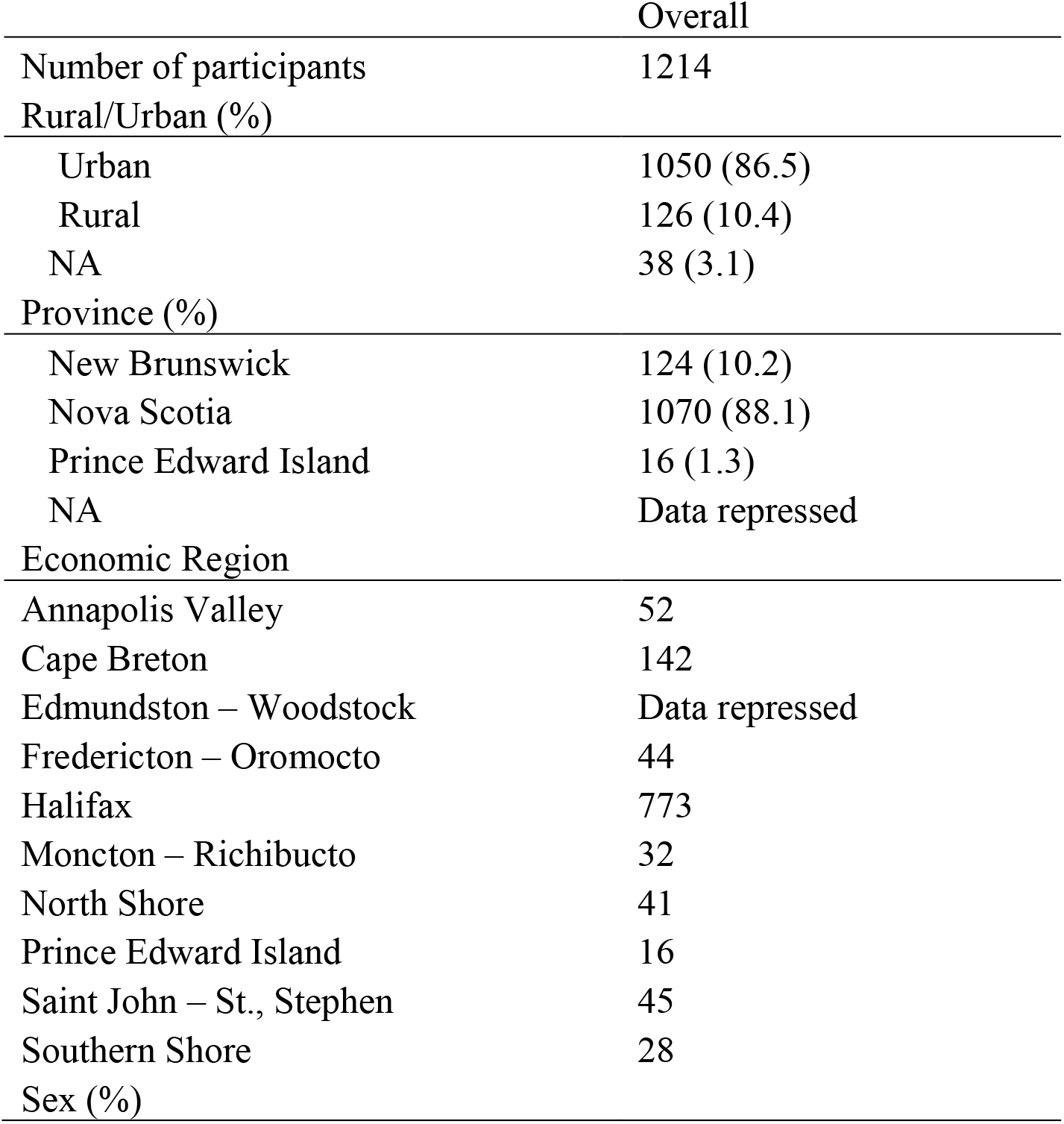
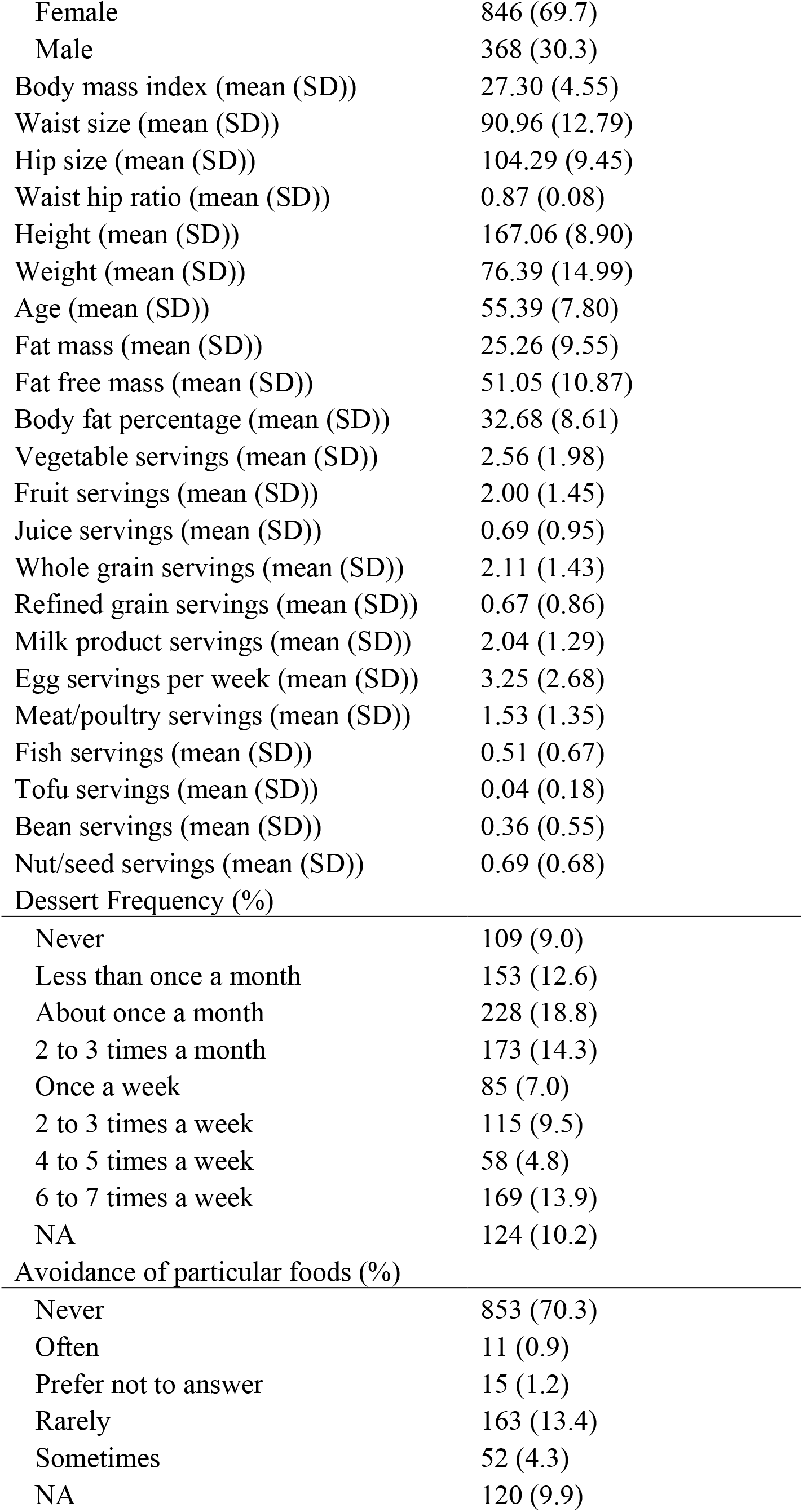
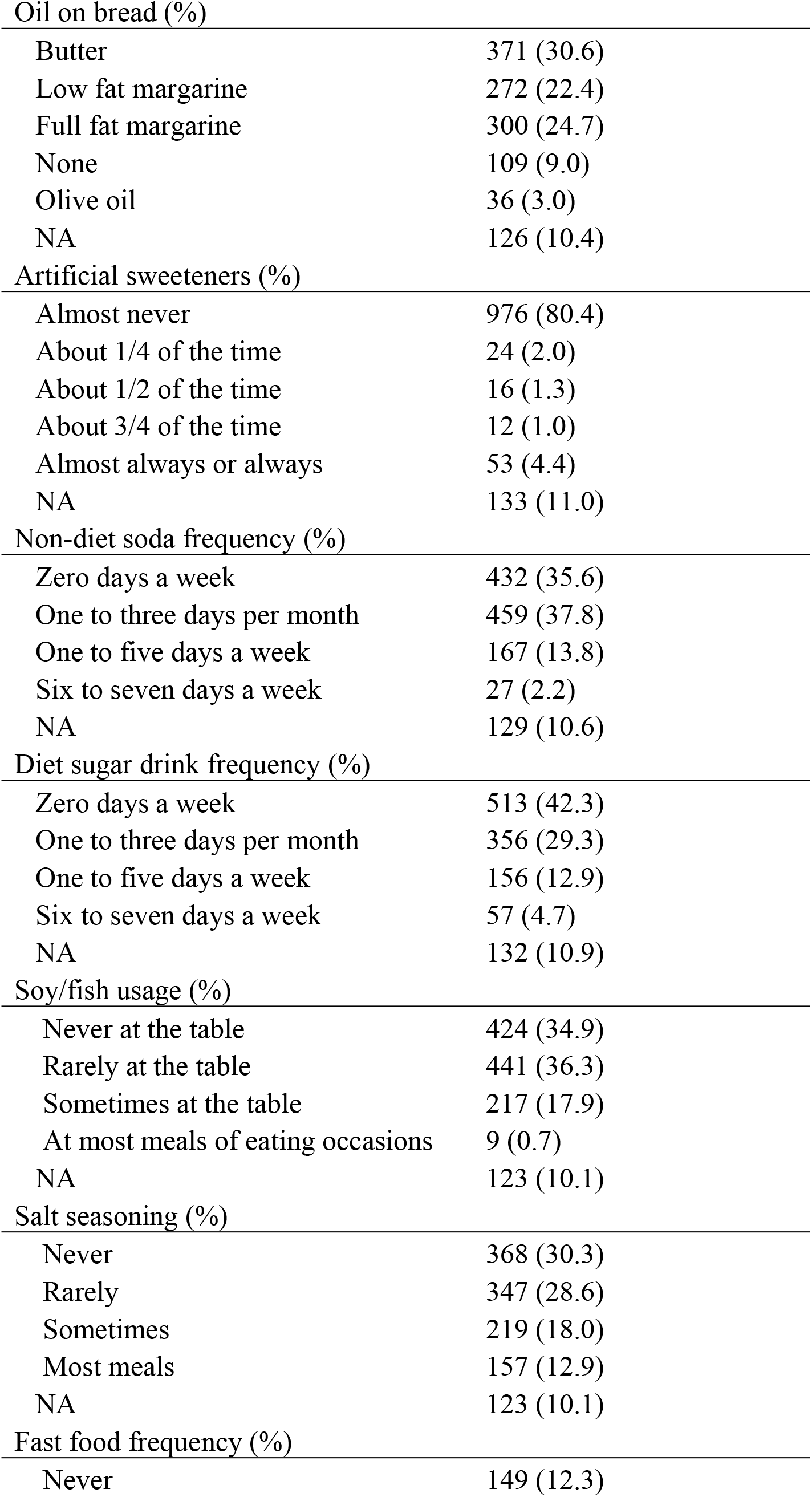
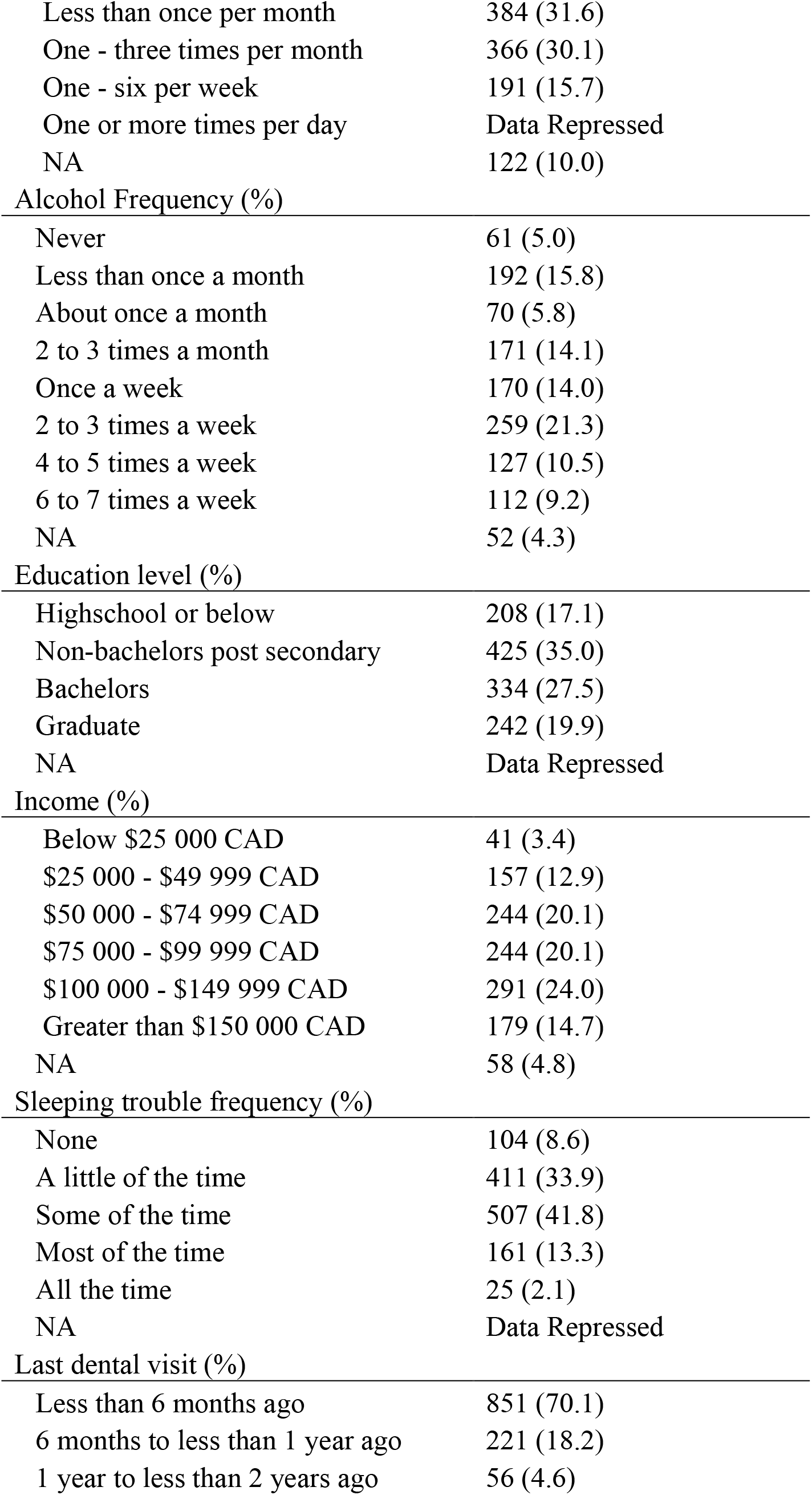
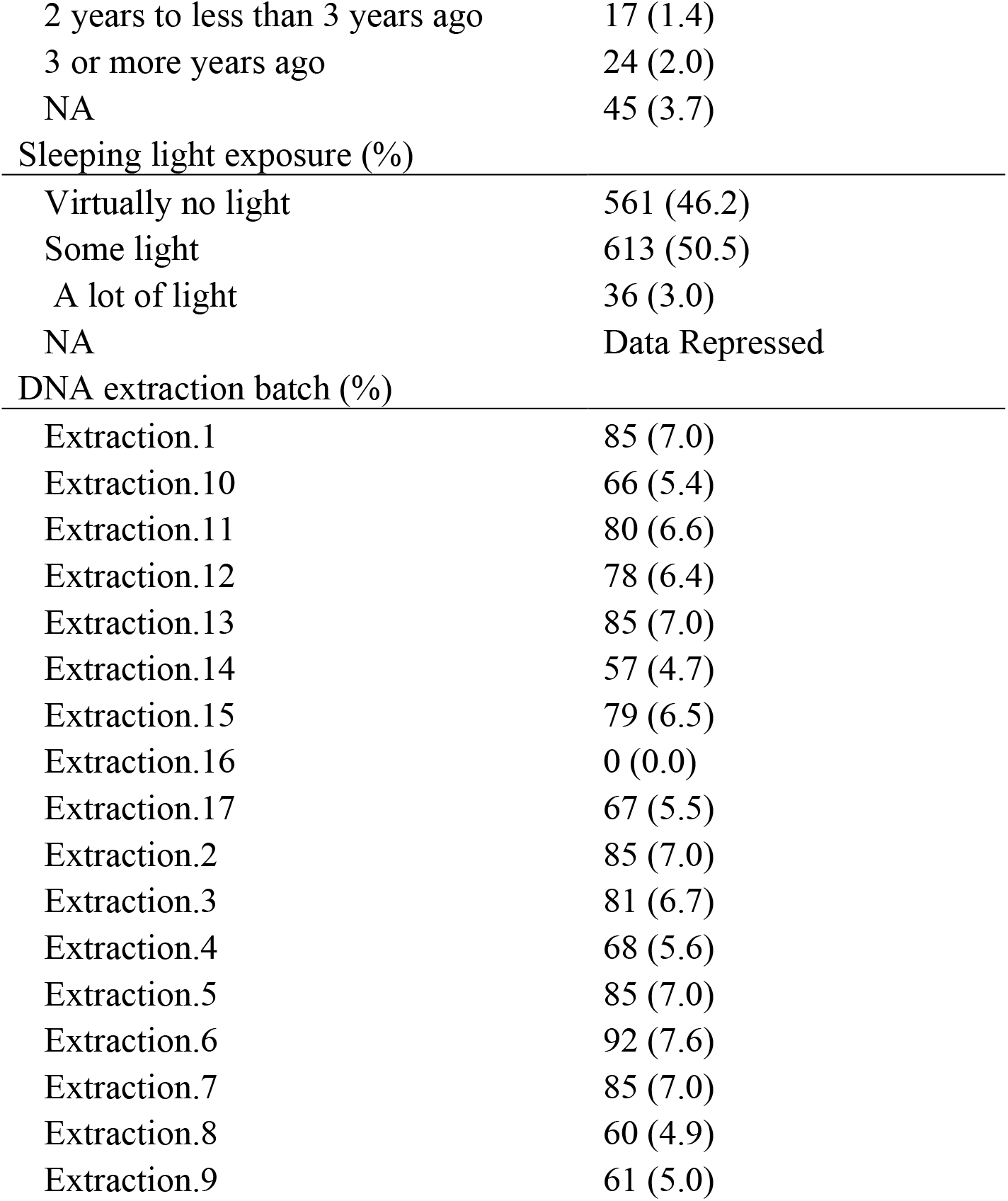
Cohort characteristic and variables compared against oral microbiome composition. NA represents responses of prefer not to answer or missing data.

|  | Overall |
| --- | --- |
| Number of participants | 1214 |
| Rural/Urban (%) |  |
| Urban | 1050 (86.5) |
| Rural | 126 (10.4) |
| NA | 38 (3.1) |
| Province (%) |  |
| New Brunswick | 124 (10.2) |
| Nova Scotia | 1070 (88.1) |
| Prince Edward Island | 16 (1.3) |
| NA | Data repressed |
| Economic Region |  |
| Annapolis Valley | 52 |
| Cape Breton | 142 |
| Edmundston – Woodstock | Data repressed |
| Fredericton – Oromocto | 44 |
| Halifax | 773 |
| Moncton – Richibucto | 32 |
| North Shore | 41 |
| Prince Edward Island | 16 |
| Saint John – St., Stephen | 45 |
| Southern Shore | 28 |
| Sex (%) |  |
| Female | 846 (69.7) |
| Male | 368 (30.3) |
| Body mass index (mean (SD)) | 27.30 (4.55) |
| Waist size (mean (SD)) | 90.96 (12.79) |
| Hip size (mean (SD)) | 104.29 (9.45) |
| Waist hip ratio (mean (SD)) | 0.87 (0.08) |
| Height (mean (SD)) | 167.06 (8.90) |
| Weight (mean (SD)) | 76.39 (14.99) |
| Age (mean (SD)) | 55.39 (7.80) |
| Fat mass (mean (SD)) | 25.26 (9.55) |
| Fat free mass (mean (SD)) | 51.05 (10.87) |
| Body fat percentage (mean (SD)) | 32.68 (8.61) |
| Vegetable servings (mean (SD)) | 2.56 (1.98) |
| Fruit servings (mean (SD)) | 2.00 (1.45) |
| Juice servings (mean (SD)) | 0.69 (0.95) |
| Whole grain servings (mean (SD)) | 2.11 (1.43) |
| Refined grain servings (mean (SD)) | 0.67 (0.86) |
| Milk product servings (mean (SD)) | 2.04 (1.29) |
| Egg servings per week (mean (SD)) | 3.25 (2.68) |
| Meat/poultry servings (mean (SD)) | 1.53 (1.35) |
| Fish servings (mean (SD)) | 0.51 (0.67) |
| Tofu servings (mean (SD)) | 0.04 (0.18) |
| Bean servings (mean (SD)) | 0.36 (0.55) |
| Nut/seed servings (mean (SD)) | 0.69 (0.68) |
| Dessert Frequency (%) |  |
| Never | 109 (9.0) |
| Less than once a month | 153 (12.6) |
| About once a month | 228 (18.8) |
| 2 to 3 times a month | 173 (14.3) |
| Once a week | 85 (7.0) |
| 2 to 3 times a week | 115 (9.5) |
| 4 to 5 times a week | 58 (4.8) |
| 6 to 7 times a week | 169 (13.9) |
| NA | 124 (10.2) |
| Avoidance of particular foods (%) |  |
| Never | 853 (70.3) |
| Often | 11 (0.9) |
| Prefer not to answer | 15 (1.2) |
| Rarely | 163 (13.4) |
| Sometimes | 52 (4.3) |
| NA | 120 (9.9) |
| Oil on bread (%) |  |
| Butter | 371 (30.6) |
| Low fat margarine | 272 (22.4) |
| Full fat margarine | 300 (24.7) |
| None | 109 (9.0) |
| Olive oil | 36 (3.0) |
| NA | 126 (10.4) |
| Artificial sweeteners (%) |  |
| Almost never | 976 (80.4) |
| About 1/4 of the time | 24 (2.0) |
| About 1/2 of the time | 16 (1.3) |
| About 3/4 of the time | 12 (1.0) |
| Almost always or always | 53 (4.4) |
| NA | 133 (11.0) |
| Non-diet soda frequency (%) |  |
| Zero days a week | 432 (35.6) |
| One to three days per month | 459 (37.8) |
| One to five days a week | 167 (13.8) |
| Six to seven days a week | 27 (2.2) |
| NA | 129 (10.6) |
| Diet sugar drink frequency (%) |  |
| Zero days a week | 513 (42.3) |
| One to three days per month | 356 (29.3) |
| One to five days a week | 156 (12.9) |
| Six to seven days a week | 57 (4.7) |
| NA | 132 (10.9) |
| Soy/fish usage (%) |  |
| Never at the table | 424 (34.9) |
| Rarely at the table | 441 (36.3) |
| Sometimes at the table | 217 (17.9) |
| At most meals of eating occasions | 9 (0.7) |
| NA | 123 (10.1) |
| Salt seasoning (%) |  |
| Never | 368 (30.3) |
| Rarely | 347 (28.6) |
| Sometimes | 219 (18.0) |
| Most meals | 157 (12.9) |
| NA | 123 (10.1) |
| Fast food frequency (%) |  |
| Never | 149 (12.3) |
| Less than once per month | 384 (31.6) |
| One - three times per month | 366 (30.1) |
| One - six per week | 191 (15.7) |
| One or more times per day | Data Repressed |
| NA | 122 (10.0) |
| Alcohol Frequency (%) |  |
| Never | 61 (5.0) |
| Less than once a month | 192 (15.8) |
| About once a month | 70 (5.8) |
| 2 to 3 times a month | 171 (14.1) |
| Once a week | 170 (14.0) |
| 2 to 3 times a week | 259 (21.3) |
| 4 to 5 times a week | 127 (10.5) |
| 6 to 7 times a week | 112 (9.2) |
| NA | 52 (4.3) |
| Education level (%) |  |
| Highschool or below | 208 (17.1) |
| Non-bachelors post secondary | 425 (35.0) |
| Bachelors | 334 (27.5) |
| Graduate | 242 (19.9) |
| NA | Data Repressed |
| Income (%) |  |
| Below \$25 000 CAD | 41 (3.4) |
| \$25 000 - \$49 999 CAD | 157 (12.9) |
| \$50 000 - \$74 999 CAD | 244 (20.1) |
| \$75 000 - \$99 999 CAD | 244 (20.1) |
| \$100 000 - \$149 999 CAD | 291 (24.0) |
| Greater than \$150 000 CAD | 179 (14.7) |
| NA | 58 (4.8) |
| Sleeping trouble frequency (%) |  |
| None | 104 (8.6) |
| A little of the time | 411 (33.9) |
| Some of the time | 507 (41.8) |
| Most of the time | 161 (13.3) |
| All the time | 25 (2.1) |
| NA | Data Repressed |
| Last dental visit (%) |  |
| Less than 6 months ago | 851 (70.1) |
| 6 months to less than 1 year ago | 221 (18.2) |
| 1 year to less than 2 years ago | 56 (4.6) |
| 2 years to less than 3 years ago | 17 (1.4) |
| 3 or more years ago | 24 (2.0) |
| NA | 45 (3.7) |
| Sleeping light exposure (%) |  |
| Virtually no light | 561 (46.2) |
| Some light | 613 (50.5) |
| A lot of light | 36 (3.0) |
| NA | Data Repressed |
| DNA extraction batch (%) |  |
| Extraction.1 | 85 (7.0) |
| Extraction.10 | 66 (5.4) |
| Extraction.11 | 80 (6.6) |
| Extraction.12 | 78 (6.4) |
| Extraction.13 | 85 (7.0) |
| Extraction.14 | 57 (4.7) |
| Extraction.15 | 79 (6.5) |
| Extraction.16 | 0 (0.0) |
| Extraction.17 | 67 (5.5) |
| Extraction.2 | 85 (7.0) |
| Extraction.3 | 81 (6.7) |
| Extraction.4 | 68 (5.6) |
| Extraction.5 | 85 (7.0) |
| Extraction.6 | 92 (7.6) |
| Extraction.7 | 85 (7.0) |
| Extraction.8 | 60 (4.9) |
| Extraction.9 | 61 (5.0) |

### Oral Microbiome 16S rRNA Sequencing

Frozen saliva samples were thawed at room temperature and aliquoted into 96 well plates. DNA from samples were then extracted using a QIAamp 96 PowerFecal QIAcube HT Kit following the manufacturer’s instructions using a TissueLyser II and the addition of Proteinase K. Sequencing of the 16S rRNA gene was performed by the Integrated Microbiome Resource at Dalhousie University. The V4-V5 region was amplified from extracted DNA in a PCR using 16S rRNA gene V4-V5 fusion primers (515FB – 926R) (32) and high-fidelity Phusion polymerase. Amplified DNA concentrations were then normalised and pooled together to be sequenced on an Illumina MiSeq. Sequencing of samples was conducted over 6 Illumina MiSeq runs producing 300 base pair paired-end reads.

#### 16S rRNA Gene Sequence Processing

Primers were removed from paired-end 300 base pair sequences using cut adapt(33). Primer free reads were then stitched together using the QIIME2 (v. QIIME2-2018.8)(34) VSEARCH(35) join-pairs plugin. Stitched reads were then filtered using the QIIME2 plugin q-score-joined using the default parameters. Quality filtered reads were then input into the QIIME2 plugin Deblur(36) to produce amplicon sequence variants (ASV). A trim length of 360 base pairs and a minimum number of reads required to pass filtering was set to 1. Amplicon sequence variants that were found in an abundance of less than 0.1% of the mean sample depth (18) were then removed from analysis. This is to keep inline with the approximate bleed-through rate on an Illumina MiSeq sequencer. After filtering a total of 13248 ASVs were recovered. Representative sequences were then placed into the Greengenes 13_8 99%(37) reference 16S rRNA tree using the QIIME2 (2019.7) fragment-insertion SEPP(38, 39) plugin. Rarefaction curves were then generated using the QIIME2 alpha-rarefaction plugin and a suitable rarefaction depth of 5000 was chosen for diversity analysis based on when the number of newly discovered ASVs came to a plateau (**Sup Fig 2**). Representative sequences were then assigned taxonomy using a custom trained V4-V5 16S rRNA naive Bayesian QIIME2 classifier(40) trained on the 99% Silva V132 database(41).

### Oral Microbiome Composition Analysis

Taxonomic composition tables were generated using the QIIME2 taxa plugin and collapsed at the genus level. All samples over 5000 reads in depth (1049) were subsampled to a depth of 5000 reads each and taxa that contributed less than a mean relative abundance of 1% were grouped together under an “Other” category. The composition stacked bar chart was then generated in R using ggplot2(42) and the x-axis was order based on the PC1 weighted Unifrac coordinates of each sample.

### Core Oral Microbiome Analysis

Taxonomic tables subsampled previously at 5000 reads were collapsed at the genus and ASV level using QIIME2. Genera/ASVs were removed at varying different sample presence cut-offs and the remaining total mean relative abundance of non-filtered out genera/ASVs was then calculated.

### Oral Microbiome Alpha Diversity analysis

Alpha diversity metrics were generated using QIIME2 (v2019.7) and the previously generated tree containing both representative sequences and reference sequences. All samples were subsampled to a depth of 5000 reads. Association between four different alpha diversity metrics (Faith’s Phylogenetic Diversity, Shannon, Evenness, Number of ASVs) were then tested using general linear models while controlling for DNA extraction. A base model containing only DNA extraction as a covariate and a testing modelling containing DNA extraction and the covariate of interest were then compared using an ANOVA and p-values were recorded. Recorded p-values were then corrected for false discovery (Benjamini and Hochberg(43)) with a chosen alpha of q < 0.1.

### Oral Microbiome Beta Diversity analysis

Beta diversity metrics were generated using QIIME2 and the previously generated phylogeny. All sequences were subsampled to a depth of 5000 reads based on the plateauing stage of rarefaction plots (**Sup Fig 2**). Association between two different beta diversity metrics (weighted UniFrac distance, Bray Curtis dissimilarity) were then tested using a PERMANOVA (adonis2 function in Vegan(44)) while controlling for DNA extraction. Marginal p values were then corrected for false discovery (Benjamini and Hochberg) and an alpha value of q < 0.1 was chosen. Significant features from univariate analysis were then included in a single multivariate model that underwent backwards covariate selection, where each co-variation with the highest p-value was removed from the model until all features were found to be significant. Additional testing using adonis2 on fat free mass and height were done while controlling for both sex and DNA extraction.

### Differential abundance analysis

Differential abundance analysis was conducted using the Corncob(45) (v 0.1.0) and Phyloseq(46) R packages. A genus level taxonomic table was generated using QIIME2 (2019.7) and genera that were not found in at least 10% of samples were removed. The fifteen covariates that were found to be significantly associated to either weighted UniFrac or Bray Curtis dissimilarities were chosen for testing. Testing of each covariate was done using the “differentialtest” function in the Corncob package while controlling for differences in DNA extraction and differential variability across DNA extraction and the covariate of interest. Heatmaps were then constructed containing any genera/ASV that were significantly associated to at least one of the covariates that were tested.

### Validation analysis

A total of 308 samples had not completely answered all 41 metadata variables of interest and therefore were removed from the original analysis. This smaller cohort was used to test our previous results by removing samples during testing of each covariate that had not answered that question on the questionnaire. Both beta diversity analysis and differential abundance analysis were carried out in the same manner as previously explained except for only testing features that were previously identified as being significantly associated with that covariate/metric. Furthermore, as there was previous evidence that these features were associated with that covariate/metric, p-values were not corrected for false discovery but an alpha value of 0.05 was chosen.

## Results

### The Healthy Oral Microbiome is Stable at the Genus Level but Variable at Higher Resolutions

We examined the oral microbiome composition of the overall cohort containing 1049 healthy individuals (**Sup Fig 1**) from Atlantic Canada to understand how anthropometric, socio-demographic and dietary choices could alter oral microbiome composition. We found that 16 genera were found to have a mean relative abundance greater than 1% (**Fig 1A**) with *Veillonella* having the largest mean contribution (21.49% +− 0.38%) followed by *Neisseria* (13.04% +− 0.40%), *Streptococcus* (11.86% +− 0.26%) and *Prevotella 7* (11.55% +− 0.24%).

**Figure 1.**
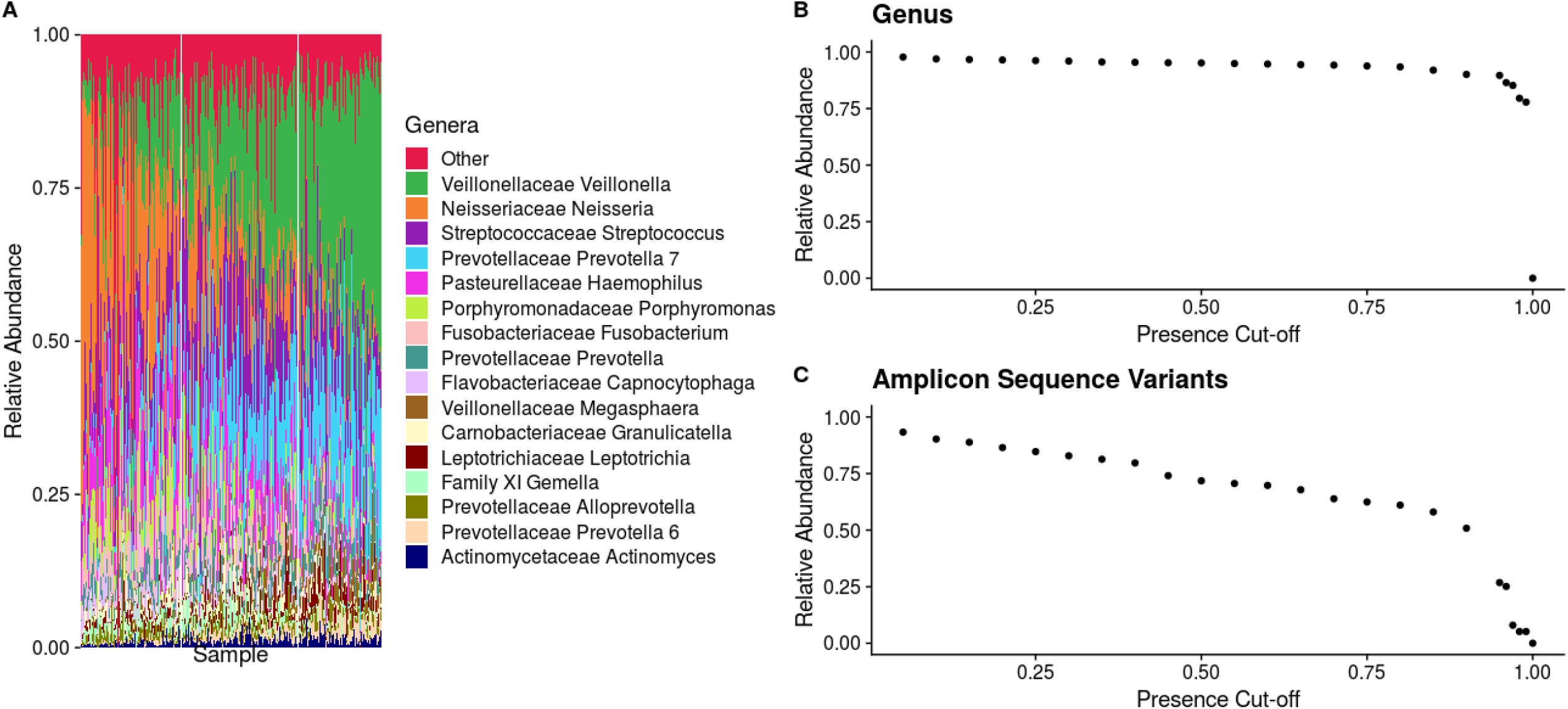
Atlantic Canadian oral microbiome composition is dominated by the genus Veillonella and is relatively similar at the genus level but highly variable at the ASV level. Samples from Atlantic Partnership for Tomorrow’s Health project (n=1049). Samples were subsampled to a depth of 5000 reads. A) Genera that had a mean relative abundance less than 1% were grouped into “Other”. B) Genera were removed at varying different sample presence cut-offs and the remaining total mean relative abundance of non-filtered genera was then calculated. C) ASVs were removed at varying different sample presence cut-offs and the remaining total mean relative abundance of non-filtered ASVs was then calculated.

To characterise the core relative abundance of core genera and ASVs within the oral microbiome of these samples the mean relative abundance of genera/ASVs that were present in greater than a specific percentage of samples was analysed. Interestingly, we found that at the genus level the oral microbiome is relatively stable with 11 genera (**Sup Fig 3A)** present in greater than 99% of all individuals making up on average a total relative abundance of 77.82% (**Fig 1B**). However, this was not the case when we examined composition at a higher taxonomic resolution. We then found that only 5.17% on average of the total relative abundance of the oral microbiome was made up of 3 ASVs (**Sup Fig 3B**) shared between 99% of all participants in the study (**Fig 1C**). These ASVs were classified as being in the *Granulicatella*, *Streptococcus*, and *Gemelli* genera but could not confidently be assigned to a specific species.

### Demographic, Anthropometric, and lifestyle choices have small but significant impacts on oral microbiome composition

We examined the relationship of both alpha and beta diversity of the oral microbiome between 41 different variables that described various demographic, lifestyle, and anthropometric measures (Table 1). Samples were split into two different cohort based on whether they had answered all 41 variables of interest. A total of 741 individuals answered all 41 variables and were included in the exploratory cohort. From this cohort we did not find any significant associations between any of the 41 variables tested and four different alpha diversity metrics (Faith’s PD, number of ASVs, Shannon, Evenness) after correction for multiple testing using linear models that were adjusted for DNA extraction batch (**Sup file 1**). We did, however, find ten variables that were associated with differences in beta diversity as measured by both weighted UniFrac (**Fig 2A**) and Bray Curtis dissimilarity (**Fig 2C**) (PERMANOVA, q < 0.1) (**Sup file 2**). We found two additional variables that were only associated with weighted UniFrac distances and three variables additional variables only associated with Bray Curtis dissimilarity (PERMANOVA, q < 0.1). Principal component analysis of both the weighted UniFrac distances and Bray Curtis dissimilarity of each sample revealed that anthropometric measures such as height, weight, waist hip ratio, waist size, and fat free mass were all correlated in similar directions along PC1, whereas features such as vegetable servings, age, and being female correlated in opposite directions (**Fig 2C)**. As sex plays an important role in determining the height, fat free mass and waist hip ratio of an individual, we attempted to determine whether sex was confounding our results from these variables. A separate analysis on weighted UniFrac distances controlling for sex indicated that fat free mass (p=0.02, r2=0.0039) and waist hip ratio (p=0.03, r2=0.0039), but not height (p=0.44, r2=0.0012) was significantly associated to microbial composition despite differences in sex. Examining the amount of variation explained by each variable by itself after controlling for DNA extraction showed small effect sizes for both weighted UniFrac distances and Bray Curtis dissimilarities (R2 0.0030 – 0.009) (**Fig 2B, 2D**). Of the features that were significant, sleeping light exposure explained the least amount of variation in both weighted UniFrac distances (r2 = 0.0036) and Bray-Curtis dissimilarity (r2=0.0030). We also found that fat free mass explained the largest amount of variation in both weighted UniFrac (r2=0.009) and Bray Curtis dissimilarity (r2=0.006). In generally we found that the rankings of effect sizes between these two different metrics agreed (Fig 2B, 2D). Also, the directionality of each feature along PC1 and PC2 were similar between both weighted UniFrac and Bray Curtis dissimilarity (Fig 2A, 2C). Examining each significant factor in our weighted UniFrac analysis using a backward selected multivariate PERMANOVA, we found that 7.0% of total oral microbiome variation could be explained by a total of 6 significant factors including DNA extraction batch despite using the same protocol, equipment and personnel for each round (**Sup Tab 1**). Interestingly, of these 6 factors DNA extraction number explained a considerable amount of the variation alone (4.18%) (**Sup Table 1**). We found similar results examining beta diversity variation using Bray Curtis dissimilarity with a slightly higher number of significant features and lower total variation explained (5.87%) (**Sup Table 2**).

**Figure 2.**
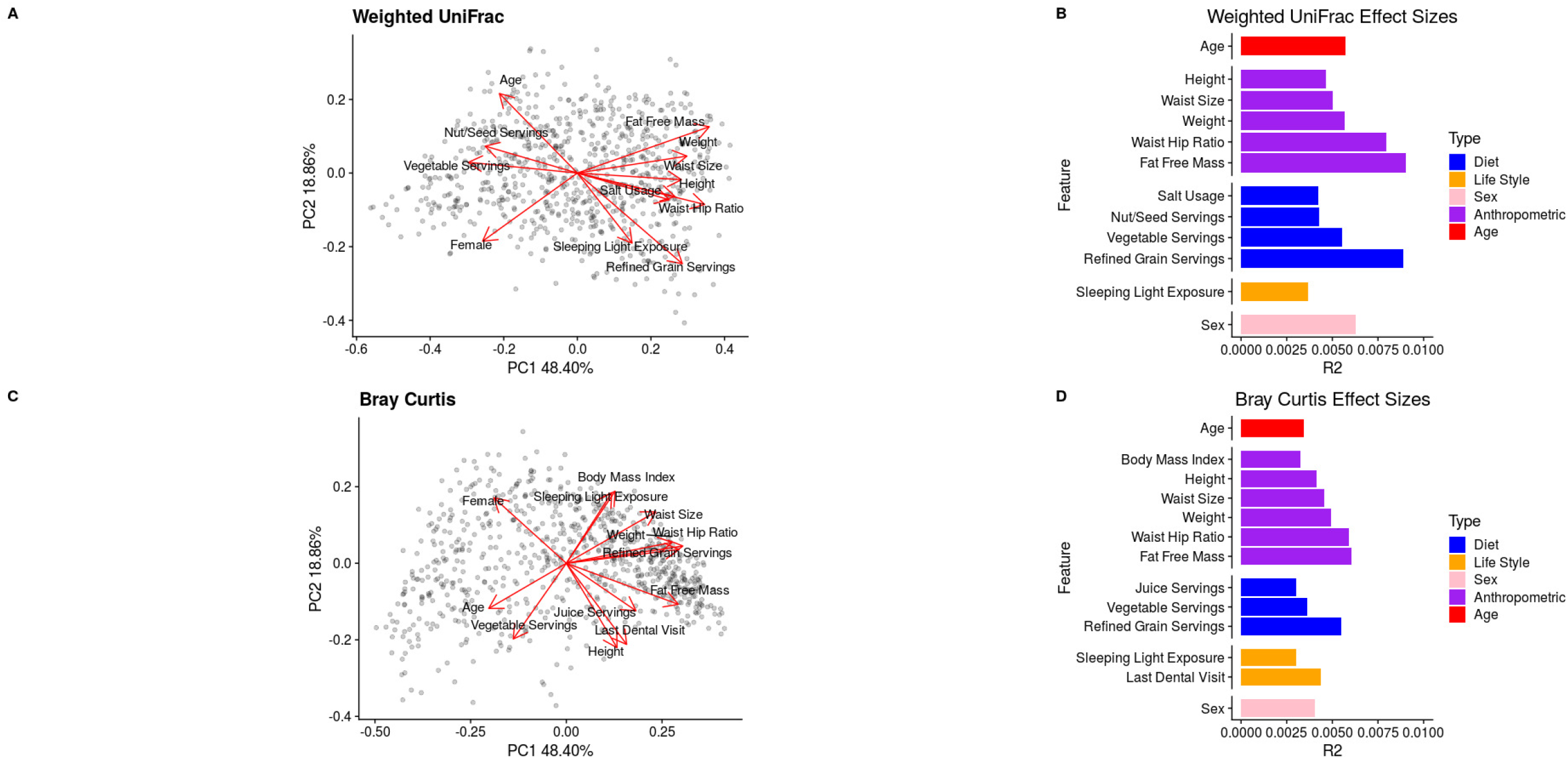
Various anthropometric, dietary and lifestyle features are significantly associated to oral microbiome composition. Saliva samples from Atlantic Partnership for Tomorrow’s Health cohort (n=741). Samples were subsampled to a depth of 5000 reads. Two different metrics were tested weighted Unifrac distances (A) and Bray-Curtis dissimilarity (C) using a PERMANOVA test while controlling for differences in DNA extraction and correction for false discovery (q < 0.1). Arrows point toward the direction each feature correlates along PC1 and PC2 while their size was scaled by the PERMANOVA R^2^ value. Panels B and D show the relative rankings of the effect sizes (R^2^) of each significant feature.

### Various oral bacterial genera and ASVs are associated with anthropometric measurements, and dietary choices in healthy individuals

We next decided to identify genera that were associated with the fifteen features previously identified as being associated with beta diversity in either the weighted UniFrac or Bray Curtis dissimilarity analysis. We found 42 genera (**Fig 3A**) and 42 ASVs (**Fig 3B**) that were significantly associated with at least one of these features after controlling for DNA extraction. We found that sex, height, and fat free mass shared similar genera and ASV associations. To control for the possibility of sex confounding our height and fat free mass associations we reanalysed the data controlling for sex. We found that no ASVs or genera were significantly associated to fat free mass after controlling for sex and only 3 genera Chloroplast, Burkholderiaceae unclassified and *Treponema 2* were significantly associated to height. Interestingly two of these three genera were not previously associated to height in our initial analysis. These results suggest that many of these features associated to height or fat free mass may be driven by differences in sex. To test this, we also tested for differences in sex while controlling for both fat free mass and height. Interestingly, we did not find any significantly associated ASVs and only three significantly associated genera Defluvittaleaceae *UCG-011*, *Leptotrichia*, and *Treponema 2*.

**Figure 3.**
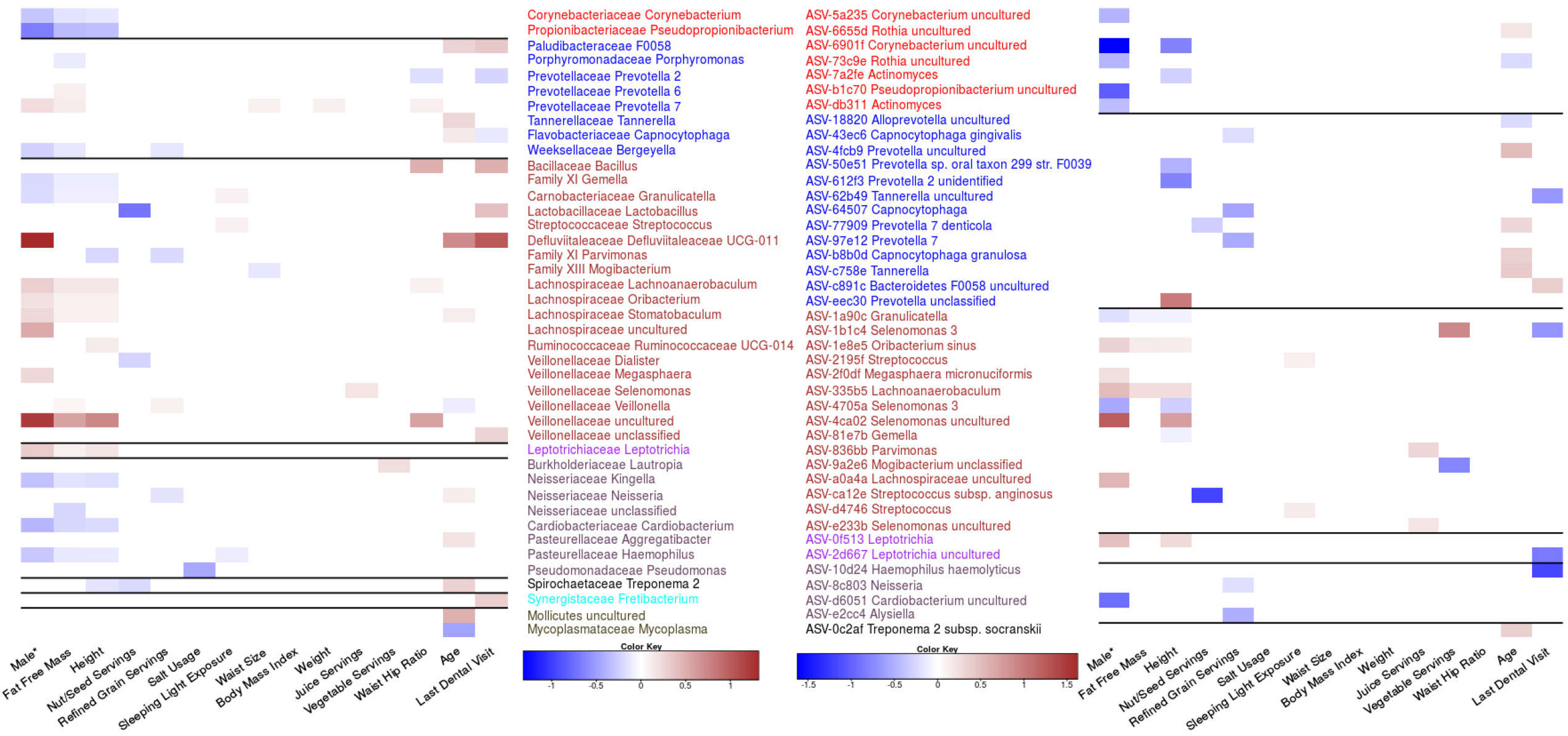
Various genera and ASVs are associated with features found to influence oral microbiome composition. Genera (A) and ASVs (B) meeting an FDR <0.1 using the Corncob R package which uses beta binomial regressions. Each feature’s false discovery rate was corrected separately, and each tested to control for differences in DNA extraction and differential variability within that feature. Ordinal variables were converted into a ranked scale for testing, and all features except for sex were scaled. *Sex was treated as a categorical value and therefore the magnitude is not directly comparable to other log odd ratios.

**Figure 4.**
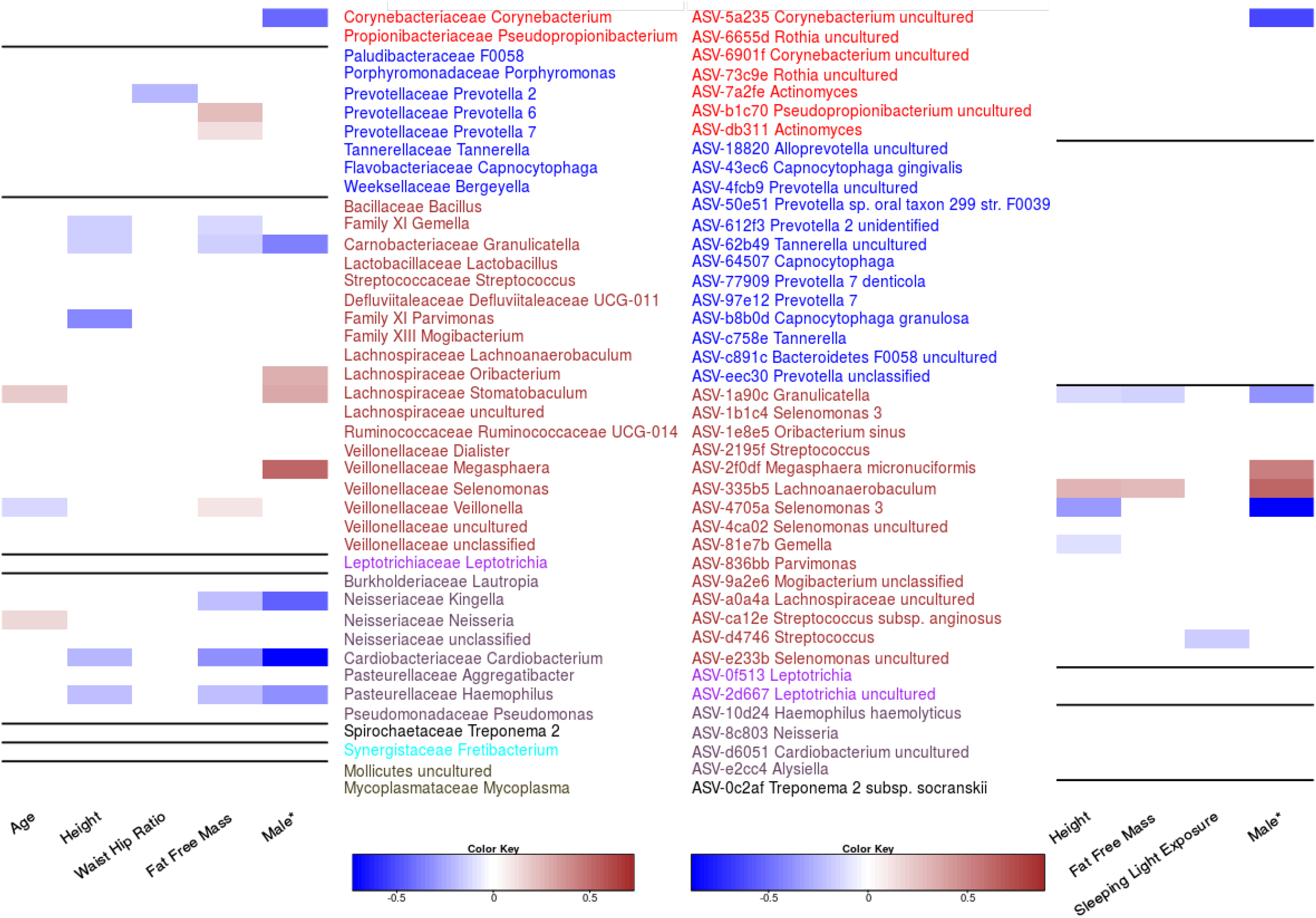
Validation of Genera and ASV association in a smaller Atlantic Partnership for Tomorrow’s Health cohort (n=308). Samples that did not complete all questions examined in this study were used to validate previous associations identified in the larger cohort. Testing procedure was done in the same manner as Figure 3 with A representing Genera and B representing ASVs.

We did not find any other features that shared similar patterns of taxonomic associations but there were multiple genera with multiple feature associations. The genus *Prevotella 7* had the highest number of features (5) associated with its relative abundance including four anthropometric measurements (height, fat free mass, waist size, waist hip ratio, and weight) and sex. Interestingly, BMI did not have any genera or ASVs significantly associated despite many other anthropometric measures showing strong taxonomic signals. We were unable to identify any single ASVs associated to waist size and weight but were able to identify a small number of genera including *Prevotella 7*, which was related to both and *Mogibacterium* with waist size. We also found that for some phyla, all taxa with significant associations had the same effect size direction. For example, genera in the Actinobacteria or Proteobacteria phyla tended to be negatively associated with fat free mass, height and being male. We also found several genera in the Proteobacteria phylum that were significantly associated with the amount of time since an individuals last dental appointment.

In contrast, examining the ASVs associated with each feature we found that in a small number of cases ASVs in the same genera had opposite directions of association to the same features. For example, two ASVs classified as *Rothia* uncultured were both significantly associated to age but in opposite directions suggesting that lower taxonomic resolution is required to identify some associations. Furthermore, we also identified cases were ASVs that were associated to a feature were classified in a genus that was found not to be related to that feature. For example, ASV-4ca02 *Selenomonas* uncultured was strongly associated with being male even though this entire collective genus was not (**Fig 3**). Further examples include ASV-e2cc4 which was classified in the genus *Alysiella*, and significantly associated with reduced refined grain servings. Examples of the opposite occurrence are also present with genera such as *Mycoplasma* being associated with age but no single ASV for this associated could be identified.

### Validation of diversity and differential abundance analysis

To help validate our findings we analyzed an additional 308 samples from a smaller subset of the Atlantic PATH cohort that had not completely answered all 41 variables of interest. We found that associations between beta diversity and anthropometric features such as height, weight, waist hip ratio, and fat free mass were recoverable within our smaller cohort (**Table 2, Sup Fig 4**). Furthermore, we also found that the associations between age and sex with oral microbiome composition were also recoverable, validating our previous analysis. We were unable to recover any significant dietary associations within this smaller validation cohort. We also were unable to recover associations between lifestyle variables such as sleeping light exposure or the time since an individuals last dental visit. The inability to recover these differences could have been due to the highly reduced sample size within this validation cohort.

**Table 2:** Validation of Beta Diversity Results

| **<u>Metric</u>** | **<u>Feature</u>** | **<u>P-value</u>** | **<u>R<sup>2</sup></u>** |
| --- | --- | --- | --- |
| **Weighted UniFrac** | Waist Hip Ratio | 0.0190 | 0.0116 |
|  | Height | 0.001 | 0.0117 |
|  | Weight | 0.010 | 0.0102 |
|  | Fat Free Mass | 0.002 | 0.0172 |
|  | Sex | 0.0390 | 0.0080 |
|  | Age | 0.0120 | 0.0105 |
| **Bray-Curtis** | Waist Hip Ratio | 0.0140 | 0.0072 |
|  | Height | 0.0030 | 0.0118 |
|  | Weight | 0.0020 | 0.0096 |
|  | Fat Free Mass | 0.0040 | 0.0110 |
|  | Waist Size | 0.0210 | 0.0065 |
|  | Age | 0.0020 | 0.0106 |
|  | Sex | 0.0380 | 0.0059 |

We further validated our differential abundance analysis using this cohort and found 8/17 genera associated with sex, 8/16 genera associated with fat free mass, 5/15 genera associated with height, and 3/11 genera associated with age were recoverable within this smaller cohort. Additionally, the negative association between *Prevotella 2* and waist hip ratio was also verified within this cohort. Furthermore, several associations between ASVs and features such as sex (5/14), height (4/12), fat free mass (2/3) and sleeping light exposure (1/2) were also found within this smaller validation cohort. All significant effect sizes that were recovered in the validation cohort except for one, between sleeping light exposure and ASV-d4746 *Streptococcus*, remained in same direction as the original cohort indicating relationships that were robust to sample choice.

## Discussion

Our analysis of 1049 healthy (**Sup Fig 1**) individuals from Atlantic Canada revealed that much of the oral microbiome of Atlantic Canadians was made up of eleven “core” genera that belong to six different phyla (*Actinobacteria*, *Fusobacteria*, *Proteobacteria*, *Firmicutes*, *Bacteroidetes*, and *Fusobacteria*). Interestingly some of these core genera found in 99% of all samples were found in relatively low abundance (<2% mean abundance) indicating that bacteria within the oral microbiome can be consistently observed with minor contributions. In contrast, the composition at the ASV level had only 3 ASVs being present in 99% of samples and only contributing 5.17% of the total oral microbiome composition on average. Overall, these results indicate that individuals tend to share similar genera within the oral cavity, but the species/strains shared between individuals is highly variable. These findings are inline with previous work from the Human Microbiome project that found the oral microbiome to be relatively stable at the genus level(1).

We found that various anthropometric and lifestyle features were significantly associated with oral microbiome composition, however, they explained only a small amount of total oral microbiome variance while controlling for DNA extraction batch (5.87-7.00%). We found that fat free mass explained the highest amount of variance (0.6-0.9%) of all biological features. While this feature had many differential abundant genera and ASVs associated with it, we were unable to recover any of them after controlling for differences in sex. This could indicate that these associations could be driven by sex and not underlying fat mass, however, we were also unable to recover many relationships between sex and taxonomic abundance while controlling for fat free mass indicating that both of these factors significant confound the other. However, despite this issue there is previous evidence to suggest that some bacteria are related to differences in body size. A study in children found reduced abundance of *Veillonella*, *Prevotella*, *Selenomonas* and *Streptococcus* in obese children (47). Interestingly, in our adult population we found similar trends with members of the *Veillonella* family being positively associated with increasing fat free mass, and members of the *Provetella* genus also being linked with higher fat free mass. Another publication on the Southern Community Cohort Study found that both *Granulicatella* and *Gemella* were associated with obesity (29) which we also found within our cohort at both the genus and ASV level. One interesting result from our study was our inability to identify any genera or ASVs linked to body mass index, despite numerous relationships between anthropometric measurements being identified. These results indicate that future studies should be advised to include sex and other measurements of body composition, such as lean body mass, when looking at relationships between the microbiome and obesity.

We found two genera *Defluviitaleaceae* UCG-011 and an uncultured genus from *Veillonellaceae* which were strongly associated with being male. However, neither of these associations were recovered in our validation cohort indicating that they could either be false positives or require a larger sample size to recover. Despite this we were still able to recover eight genus level associations in our validation cohort, however, only a few of these associations match those that were previously reported. Renson et al. found two genera *Lactobacillus* and *Actinobacillus* to be higher in males, which we did not find within our study (23). This could have been due to multiple differences including sampling procedures or systemic protocol bias. Raju et al. found that there was a high relative abundance of *Haemophilus* in females which we also found in our study, however they also found *Oribacterium* to be increased in females which was opposite from what was found in this study (47). Differences between these studies and ours can in part be attributed to differences in samples collection and sequencing primers used highlighting the importance of standardizing protocols within the field.

We were unable to recover any relationships between dietary features within our validation cohort, however, refined grain servings per day had the largest impact on oral microbiome composition in our initial analysis. During this initial analysis, we found that bacteria from four genera *Bergeyella*, *Parvimonas, Veillonella,* and *Neisseria* decreased in relative abundance with increasing refined grain intake. Interestingly, refined grain intake had a very strong association with inflammatory bowel disease in a previous analysis of this cohort(48), and alterations in the oral microbiome have been linked to inflammatory bowel disease in the past (49). Previous work by Said et al., found multiple genera in differential abundance between individuals with and without IBD including the increased presence of *Veillonella* (16), which we found to be linked positively with refined grain intake.

Other dietary factors we found linked to oral microbiome composition in our original analysis include both juice servings and vegetable servings. However, were only able to find a small number of genera and taxa linked to vegetable serving intake and juice serving intake. Furthermore, we were unable to recover these effects in our validation cohort indicating the possibility for a false positive or the requirement of a large sample size to see these effects. Previous work within the field has found conflicting evidence on the role of diet impacting oral microbiome composition. This previous evidence along with the inability to recover these relationships within our validation cohort indicates that diet may only have a small impact on oral microbiome variation, and that these effects require large samples to recover them or that different dietary capture methods have a strong influence on the observed results.

Looking at all features that were significantly associated to oral microbiome composition together in a single model we were only able to explain a small portion of the total variance between samples (6.75-6.92%). This indicates that while many of these features are significantly related to microbial composition each one by themselves tends to only cause small shifts in overall microbial composition. Furthermore, a majority of the variance accounted for was due to differences in DNA extraction date. This shows that while slight technical variations such as the time when DNA extraction was done can have larger impacts on sample composition emphasizing the need to control for these technical variations during large population-based studies.

One large limitation to our study was our lack of detailed dental history information from participants. While we did record how recently each individual last visited the dentist, we were unable to retrieve detailed information on dental health, which has been found to have dramatic impacts on oral microbiome composition (19). This could explain some of the missing variation that was not accounted for in our study, however, it is unlikely to explain all 93.25% indicating we are still missing a suitable amount of information on what determines an individual’s oral microbiome composition.

In conclusion, our study indicates that the healthy oral microbiome is relatively stable between individuals at the genus level and is impacted very little by any one factor. Future studies that attempt to identify oral microbial biomarkers associated with disease may be encouraged by the lack of major confounding variables and may be justified in controlling only for sex, body composition, oral health, and basic dietary information.

## Availability of data and materials

All sequencing data has been uploaded to the European Nucleotide Archive and is available under the accession number PRJEB38175. Code used to analysis all data is available at https://github.com/nearinj/Nearing_et_al_2020_Oral_Microbiome. De-identified metadata used in this project can be accessed by contacting the Atlantic Partnership for Tomorrow’s Health project.

## Funding

JTN is supported by both a Research Nova Scotia, Scotia Scholars award (2019-2022) as well as a Nova Scotia Graduate Scholarship (2019-2023). JVL was supported by a Canadian Institutes of Health Research (CIHR)-Canadian Association of Gastroenterology-Crohn’s Colitis Canada New Investigator Award (2015–2019), a Canada Research Chair Tier 2 in Translational Microbiomics (2018-2019) and a Canadian Foundation of Innovation John R. Evans Leadership fund (awards #35235 and #36764), a Nova Scotia Health Research Foundation (NSHRF) establishment award (2015–2019), an IWK Health Centre Research Associateship, a Future Leaders in IBD project grant, a donation from the MacLeod family and by a CIHR-SPOR-Chronic Diseases grant (Inflammation, Microbiome, and Alimentation: Gastro-Intestinal and Neuropsychiatric Effects: the IMAGINE-SPOR chronic disease network). The data used in this research were made available by the Atlantic Partnership for Tomorrow’s Health (Atlantic PATH) study, which is the Atlantic Canada regional component of the Canadian Partnership for Tomorrow’s Health Project funded by the Canadian Partnership Against Cancer and Health Canada. The views expressed herein represent the views of the authors and do not necessarily represent the views of Health Canada.

## Acknowledgements

This research has been conducted using Atlantic PATH data and biosamples, under application #2018-103. We would like to thank the Atlantic PATH participants who donated their time, personal health history and biological samples to this project. We would also like to thank the Atlantic PATH team members for data collection and management.

